## Supplementary material for "LucID: A Self-Activating Bioluminescent Biotin Ligase for Light-Free Proximity Labeling in Deep Tissues": https://doi.org/10.5281/zenodo.15631269: calc_sym_rmsd.py

```
from pyrosetta import *
init()
pose = pose_from_file("sym_complex.pdb")
reference_pose = pose_from_file("reference.pdb")
rmsd = CA_rmsd_symmetric(pose, reference_pose)
print(f"Symmetric RMSD: {rmsd:.2f} Å")
```
