## Supplementary material for "LucID: A Self-Activating Bioluminescent Biotin Ligase for Light-Free Proximity Labeling in Deep Tissues": https://doi.org/10.5281/zenodo.15631269: README.txt

LucID Computational Supplement
==============================

This package contains all computational protocols, configurations, and scripts 
used for modeling and validating the LucID protein system as described in the manuscript:
"LucID: A Self-Activating Bioluminescent Biotin Ligase for Light-Free Proximity Labeling in Deep Tissues"

Directory Structure
-------------------
.
├── Protocols/               # RosettaScripts XML files for Relaxation, QM/MM, Docking
│   ├── relax_protocol.xml
│   ├── custom_relax.xml
│   ├── qmmm.xml
│   ├── symmetric_dock.xml
│   └── symmetric_qmmm.xml (optional)
├── Configs/                 # Configuration files for QM and symmetry
│   ├── ref2015.wts
│   ├── xtb_config
│   ├── lucid.span
│   ├── C3.symm
│   └── cys.cst
├── Scripts/                 # Python scripts for result validation
│   ├── check_qm.py
│   └── calc_sym_rmsd.py
├── Templates/               # Gaussian input and active-site PDB for QM
│   ├── QM_region.pdb
│   └── gaussian_template.com
├── Data/
│   ├── residue_energies.csv
│   ├── lucid_scores.sc
│   └── mpi_flags
├── SLURM/
│   └── rosetta_slurm.sh

Usage Overview
--------------
1. **Relaxation (Standard)**
   Command:
     relax.linuxgccrelease @mpi_flags

2. **Membrane/Disulfide Relaxation**
   Command:
     rosetta_scripts.linuxgccrelease -parser:protocol custom_relax.xml -s lucid_initial.pdb

3. **Symmetric Docking**
   Command:
     rosetta_scripts.linuxgccrelease -parser:protocol symmetric_dock.xml -s lucid.pdb

4. **QM/MM Activation Mechanism**
   Command:
     rosetta_scripts.linuxgccrelease -parser:protocol qmmm.xml -s top_model.pdb -qmmm:config xtb_config

5. **Energy Landscape Analysis**
   Use: `check_qm.py`, `calc_sym_rmsd.py` for validating QM convergence and symmetry

Dependencies
------------
- Rosetta (3.13+)
- Python 3.7+ with pandas, seaborn, matplotlib
- xtb or Gaussian for QM/MM (optional)
- SLURM or MPI (for cluster-based parallel runs)

Notes
-----
- The provided `.wts` file customizes the ref2015 energy function
- `lucid.span` defines transmembrane regions (used with membrane protocol)
- `cys.cst` constrains disulfide bonds (res 23-128)
- All files are compatible with standard Rosetta tools
- Files were manually curated for publication reproducibility

Author: Afsaneh Taheri Kal-Koshvandi
Contact:

License: For research use only. Not for redistribution without permission.
