## Supplementary material for "LucID: A Self-Activating Bioluminescent Biotin Ligase for Light-Free Proximity Labeling in Deep Tissues": https://doi.org/10.5281/zenodo.15631269: rosetta_slurm.sh

```
#!/bin/bash
#SBATCH --job-name=LucID_relax
#SBATCH --nodes=4
#SBATCH --ntasks-per-node=12
#SBATCH --cpus-per-task=1
#SBATCH --mem=16G
#SBATCH --time=24:00:00
#SBATCH --output=rosetta_%j.out
#SBATCH --partition=gpu

module load openmpi/4.1.1
module load rosetta/3.13

mpirun -np 48 relax.mpi.linuxgccrelease @mpi_flags
```
