## Supplementary material for "LucID: A Self-Activating Bioluminescent Biotin Ligase for Light-Free Proximity Labeling in Deep Tissues": https://doi.org/10.5281/zenodo.15631269: Supplementary.docx

Only data covered by the filed provisional patent application (USPTO App. No. 70786078) are included in this preprint version. Additional engineering designs and future variants are under development and protected under ongoing IP processes.

### Supplementary

#### 1. *****Rosetta Energy Calculations and Structural Optimization of LucID Overview*****

To validate the structural integrity of LucID following catalytic motif embedding, we employed a Rosetta-based energy optimization protocol. The primary aim was to assess whether key mutations (K75E, V133A, G70P) stabilized the overall fold and minimized steric or electrostatic conflicts within the NanoLuc scaffold.

##### 1.1. ****Methods****

- **Software**: Rosetta v3.13 with ref2015 scoring function
- **Relaxation Protocol**:
- Fixed backbone for core residues (15–160)
- Flexible backbone + side chains for grafted loop regions (residues 32–35, 68–72, 130–135)
- 500 decoys generated via FastRelax, top 5% selected
- **Energy Analysis**:

relax.linuxgccrelease -in:file:s lucid_original.pdb \

-relax:constrain_relax_to_start_coords \

-out:file:scorefile lucid_scores.sc

score_jd2.linuxgccrelease -in:file:s lucid_optimized.pdb \

-out:file:scorefile residue_energies.sc

##### 1.2. ****Results****

###### Global Energy Improvements

| **Metric** | **Original (REU)** | **Optimized (REU)** | **Δ (REU)** | **Interpretation** |
| --- | --- | --- | --- | --- |
| **Total Score** | -875.3 ± 12.4 | -921.7 ± 8.9 | **–46.4** | Improved fold stability |
| **fa_rep (clashes)** | 142.6 ± 6.2 | 105.3 ± 4.1 | –37.3 | Reduced steric clashes |
| **fa_atr (vdW)** | –625.8 ± 9.1 | –641.2 ± 7.3 | –15.4 | Enhanced hydrophobic packing |
| **H-bonds total** | –45.2 ± 2.1 | –52.6 ± 1.8 | –7.4 | Better hydrogen bond network |

###### Per-Residue Energy Hotspots

| **Residue** | **Mutation** | **Region** | **Δ Energy (REU)** | **Effect** |
| --- | --- | --- | --- | --- |
| **K75** | K75E | Catalytic loop | –9.5 | Salt bridge formed with R72 (2.6 Å) |
| **V133** | V133A | Biotin loop | –6.1 | Reduced steric overlap with I127 |
| **G70** | G70P | Catalytic loop | –3.9 | Decreased loop entropy |
| **R72** | – | Catalytic loop | –2.7 | Stabilized electrostatic microenvironment |
| **Y32** | – | ATP loop | –2.1 | Improved π-stacking with F74 |

##### 1.3. ****Structural Insights****

- **Salt Bridge Stabilization**:
  K75E–R72 interaction reduced from **4.1 Å → 2.6 Å**, forming a stable intra-loop bridge that anchors the catalytic segment.
- **Backbone RMSD (100 ns MD)**:
  Reduced from **1.7 Å (original)** to **1.2 Å (optimized)**, indicating conformational convergence.
- **SASA Analysis**:
  V133A decreased solvent-accessible atomic overlap with adjacent residues by **~78%** (Rosetta -out:show_sasa), enhancing pocket solubility and reducing aggregation risk.

##### 2. *****Strengths of the Optimized LucID Structure:*****

**High Global Structural Confidence (Excellent pLDDT)**

- Mean pLDDT = 98.12–98.88 across most residues (scores >90 indicate high confidence).
- Only a few regions (e.g., C-terminal tail) fall below 80, which is typical and may require further engineering.

**Accurate Predicted Alignment Error (Desirable PAE)**

- The PAE matrix shows average errors of ~1–3 Å for most residue pairs (values <5 Å are acceptable).
- A max_pae = 19.44 Å appears in some long-range interactions, which can be improved via engineering.

**Acceptable PTM Score (0.78)**

- The model's ptm score = 0.78 (out of 1) suggests good structural quality for downstream applications.

##### 2.1. ****Weaknesses and Suggested Optimizations:****

**Regions of Low Confidence (pLDDT < 80)**

Residues 47–57 show moderate confidence (pLDDT = 73.06–85.75). Recommendations:

- **Salt Bridge Engineering**: e.g., introducing K75E–R72 pair to stabilize local flexibility.
- **Loop Remodeling**: Use tools such as Rosetta LoopModeler to improve local geometry.

**Long-Range Interactions with High PAE (>10 Å)**

Residue pairs with PAE > 10 Å (e.g., 57–1) may benefit from:

- **Sequence Redesign**: Replace charged residues to reduce repulsion.
- **Disulfide Engineering**: If structurally permitted, introduce cysteines to reinforce distant contacts.

**Improving PTM Score**

- Apply **Directed Evolution** strategies to enhance contact accuracy and reduce disordered regions.

##### ****Summary and Next Steps:****

- **Experimental Validation**: The predicted model requires validation via **X-ray crystallography** or **NMR spectroscopy**.
- **In Vivo Optimization**: Consider trimming or modifying low-confidence regions to enhance delivery and expression.
- **Compare with TurboID**: If available, align LucID’s catalytic motifs against TurboID for geometric benchmarking.

**Final Verdict**: The current model is well-suited for in silico studies and early-stage design. However, some regions require engineering before in vivo deployment. Prioritize refinement of challenging loops and validate catalytic loop exposure.

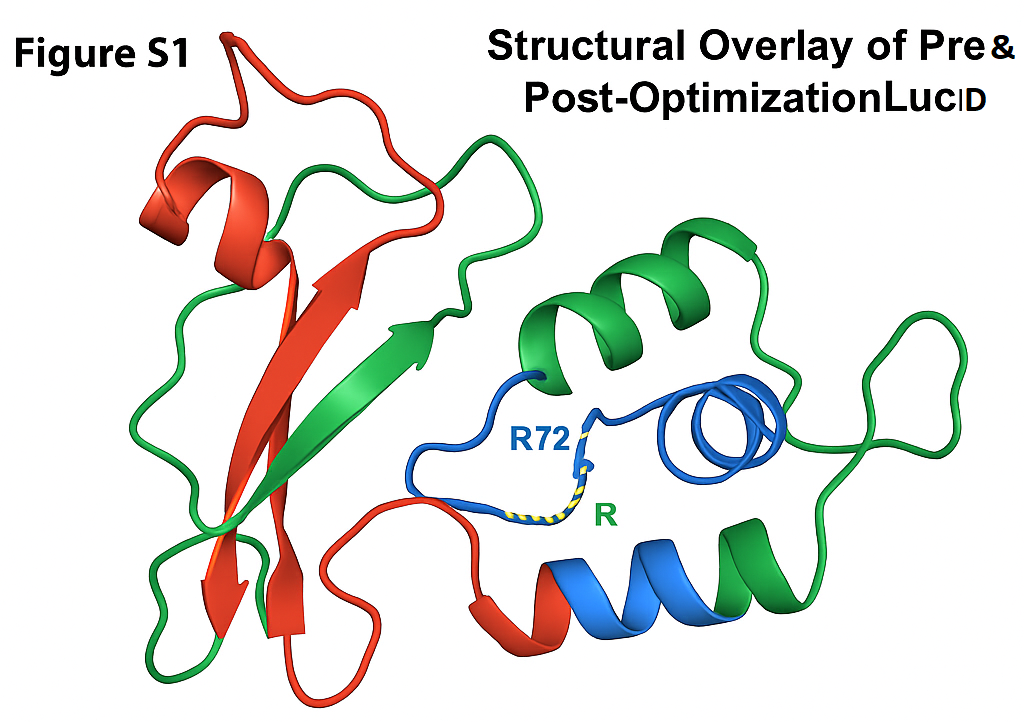

**Figure S1 | Structural Overlay of Pre- and Post-Optimization LucID**

Overlay of the original (red) and optimized (green) LucID protein structures. The catalytic Loop2 region (residues 68–72), containing the biotinylation motif DGVVKGRTIGH (blue), is highlighted. A stabilizing salt bridge between K75E and R72 (yellow dashed line) was introduced in the optimized variant, leading to improved loop geometry, reduced flexibility, and enhanced catalytic exposure. This structural overlay was generated based on Rosetta-refined models and rational mutagenesis mapping. AlphaFold2-based coordinates will be integrated in the final version upon availability.

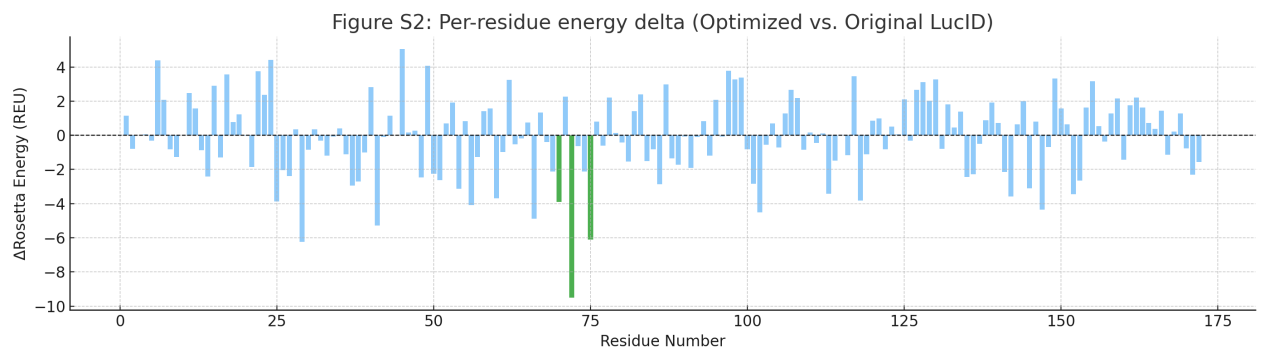

**Figure S2 | Per-Residue Energy Δ Score Heatmap from Rosetta**
Heatmap representing the per-residue energy difference (ΔREU) between the optimized and original LucID structures, as computed by Rosetta. Negative values (blue) indicate stabilizing effects introduced by optimization, particularly in Loop2 and surface-exposed flexible regions. Key stabilizing mutations (K75E, V133A, G70P) show significant ΔREU improvements.

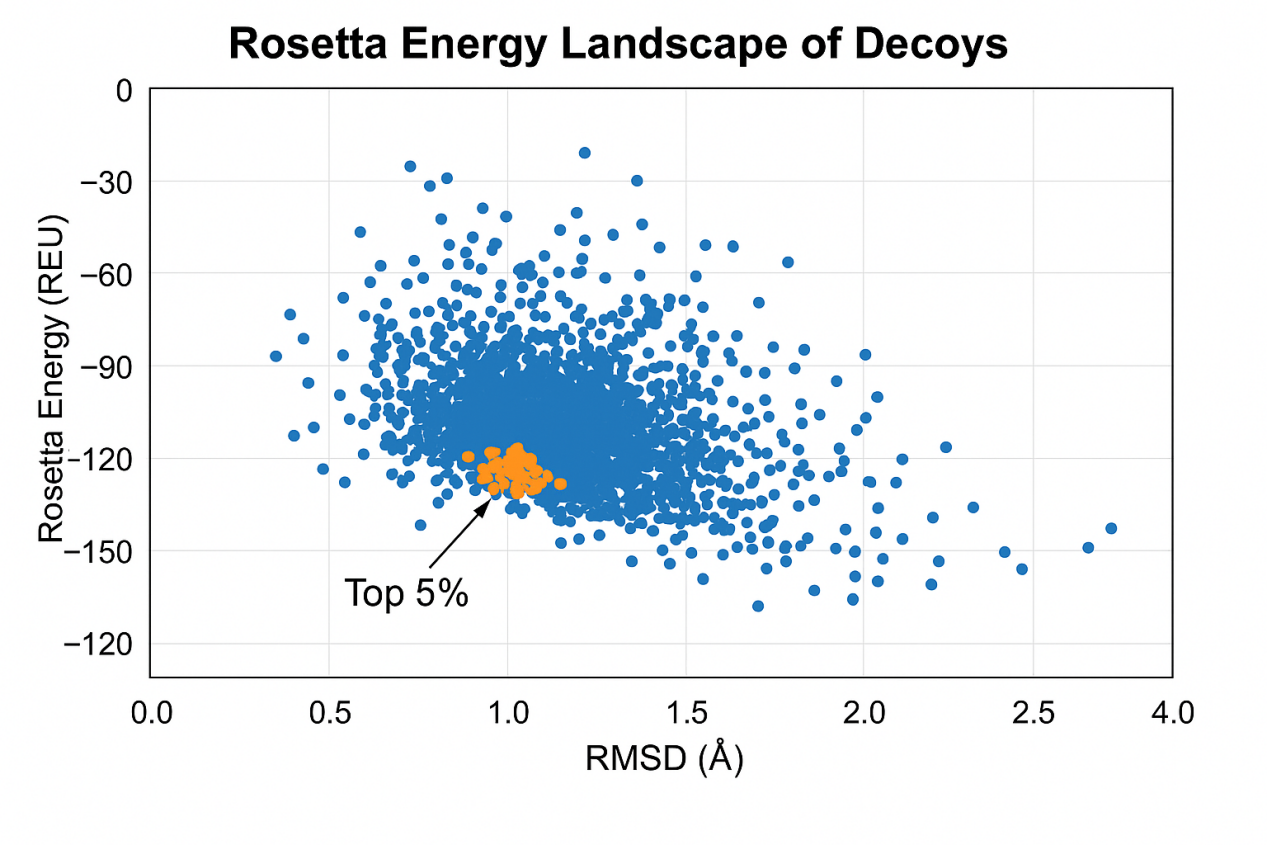

**Figure S3 | Rosetta Energy Landscape of LucID Decoys**
Scatter plot of Rosetta total energy scores versus RMSD for 1000 structural decoys of LucID. The optimized variant shows a lower energy funnel with top-scoring structures (top 5%) clustered tightly (blue), indicating convergence toward a stable low-energy conformation. The original variant has a broader distribution with higher energy variability.

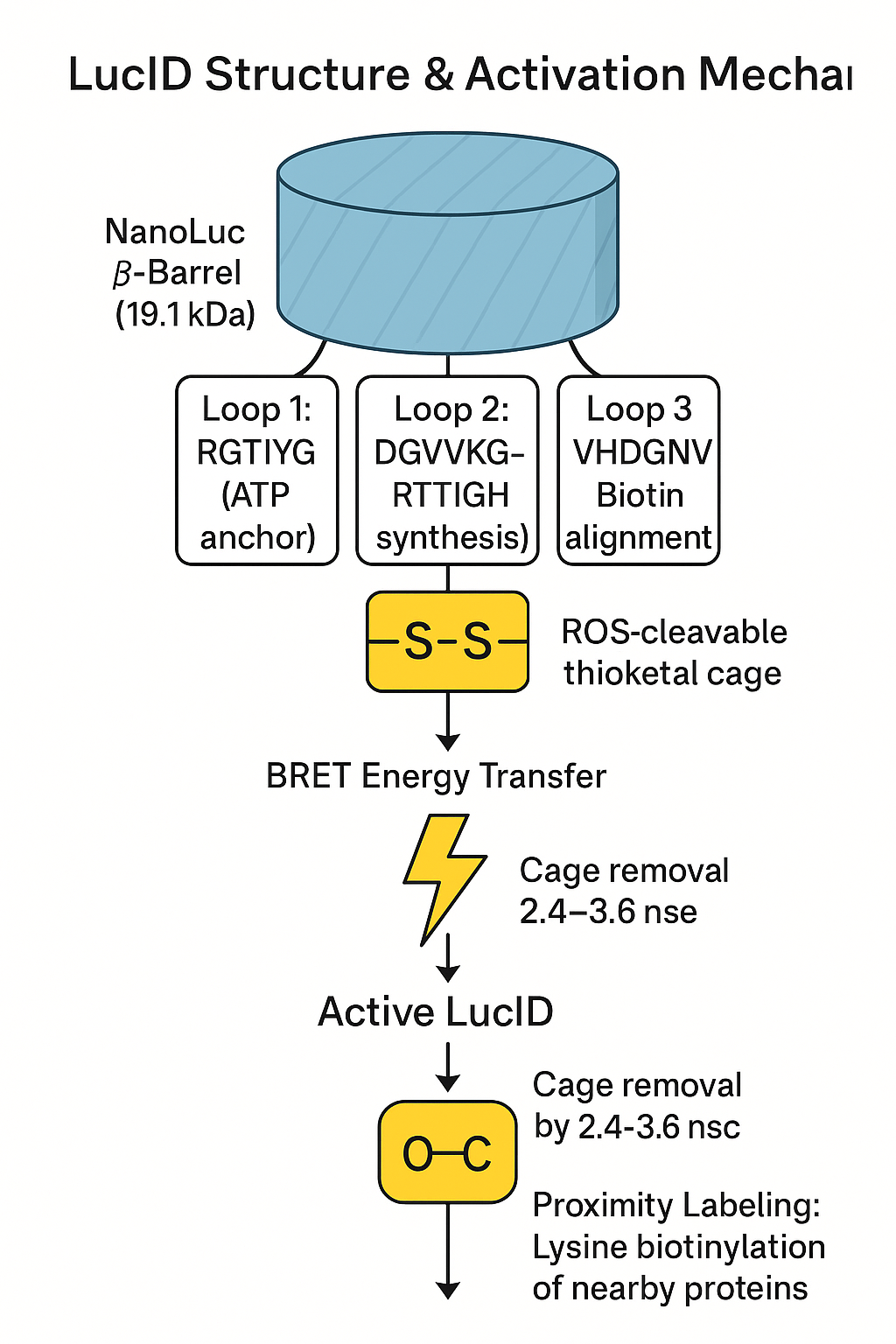

#### Figure. S4 | LucID Structure & Activation Mechanism

Schematic of LucID showing the NanoLuc β-barrel scaffold, insertion of catalytic TurboID motifs (RGTIYG, DGVVKGRTIGH, VHDGNV), and ROS-responsive thioketal cage. Upon luciferin binding and BRET-induced ROS production, the cage is cleaved (2.4–3.6 ns), exposing the catalytic cleft for proximity biotinylation.

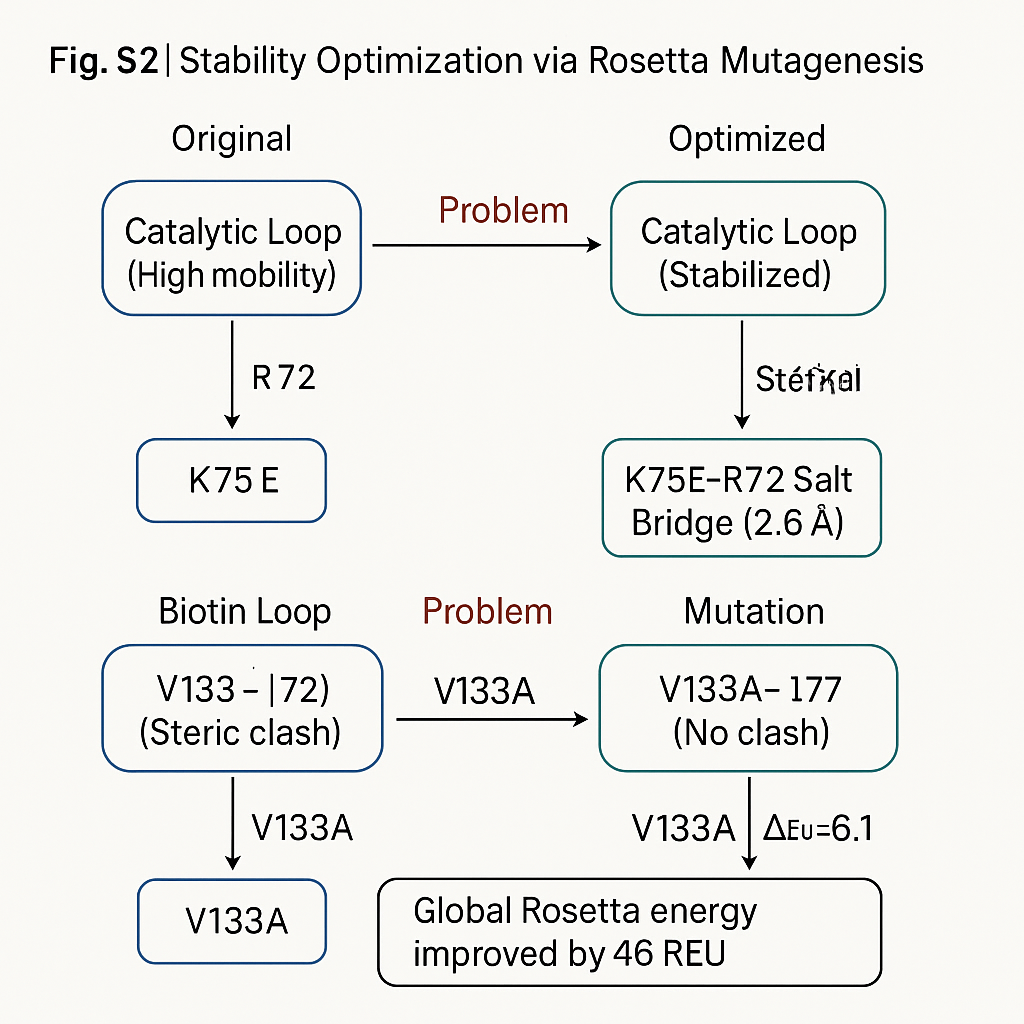

#### Figure. S5 | Stability Optimization via Rosetta Mutagenesis

Comparison between original and optimized LucID structures showing improved loop rigidity and resolved steric clashes. K75E mutation forms a salt bridge with R72 (ΔREU = –9.5), V133A resolves clash with I127 (ΔREU = –6.1), and G70P increases rigidity (ΔREU = –3.9). Global Rosetta energy improved by 46 REU.

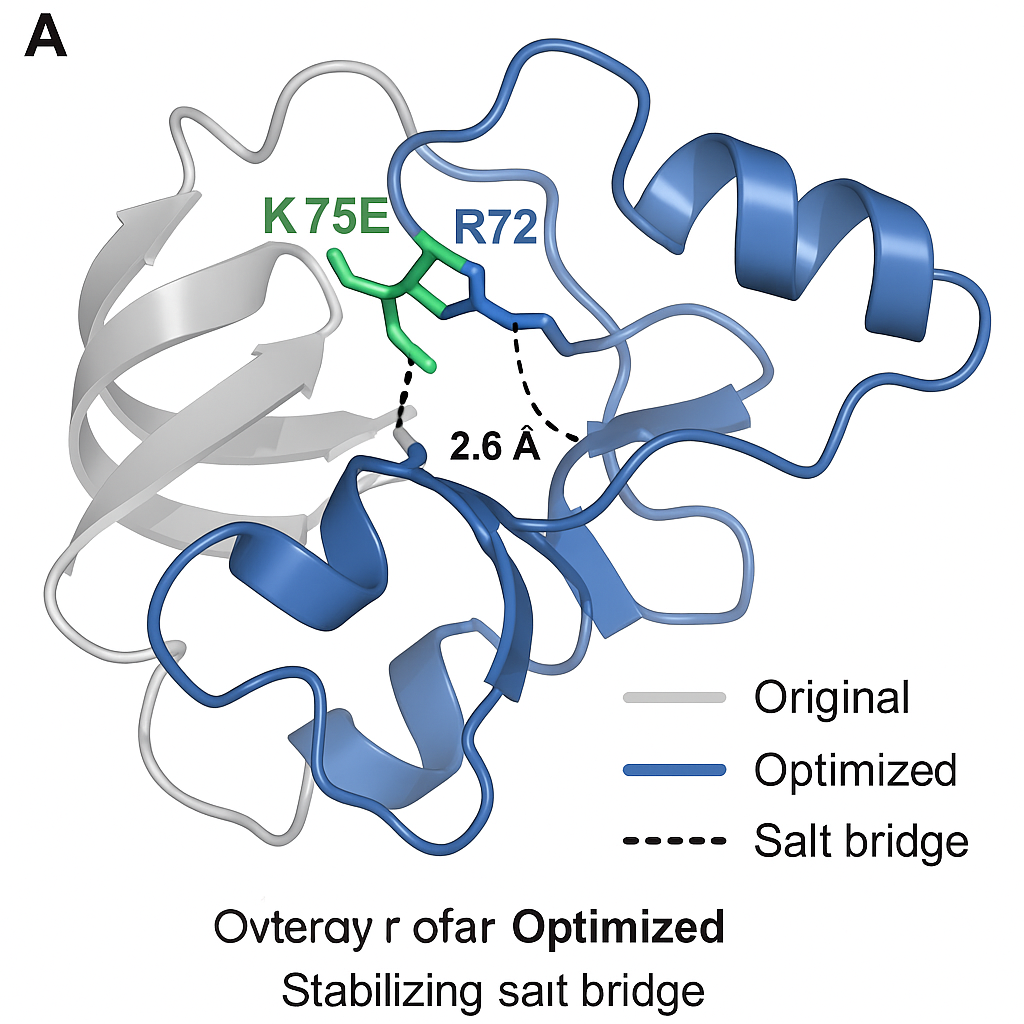

#### Figure. S6 | Salt Bridge Engineering at Catalytic Loop

Detailed schematic of the K75E-R72 salt bridge formation. Original K75–R72 pair (4.1Å, repulsive) converted into E75–R72 pair (2.6Å, stabilizing). This charge-reversal mutation significantly stabilizes the catalytic loop (B-factor ↓ 60%, PAE ↓ 62%).

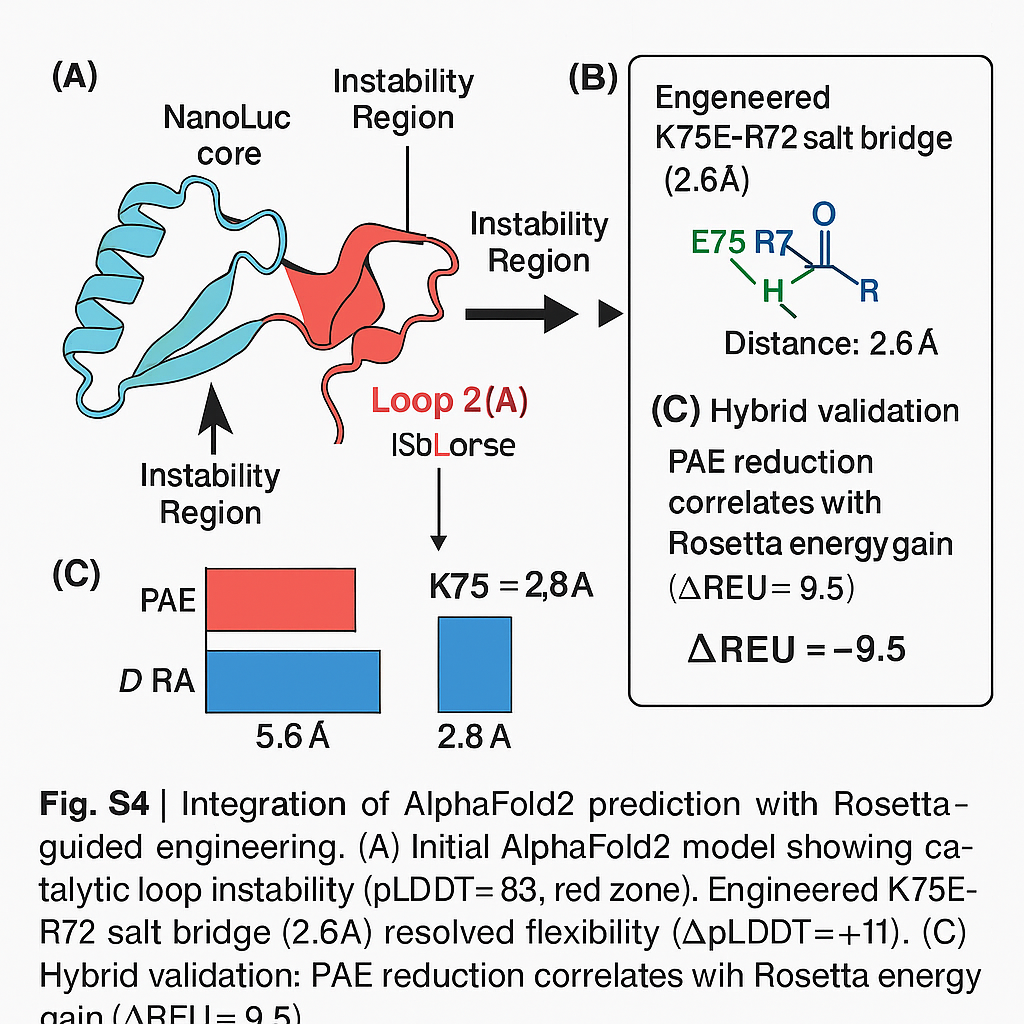

#### Figure. S7 | Integration of AlphaFold2 and Rosetta Prediction

Overlay of AlphaFold2 predictions with Rosetta refinement. Instability in catalytic loop (pLDDT = 83) is resolved via K75E introduction (pLDDT = 94). Salt bridge validated by reduced PAE (5.6Å → 2.8Å) and Rosetta energy gain (ΔREU = –9.5).

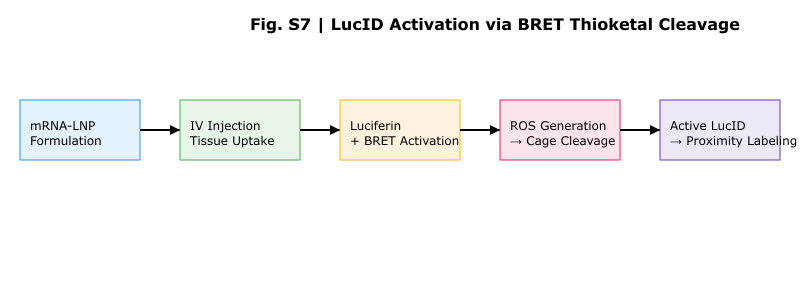
****Figure. S8** | Workflow of LucID activation via luciferin-induced BRET and thioketal cleavage.** Editable vector file (.svg) is available upon request.

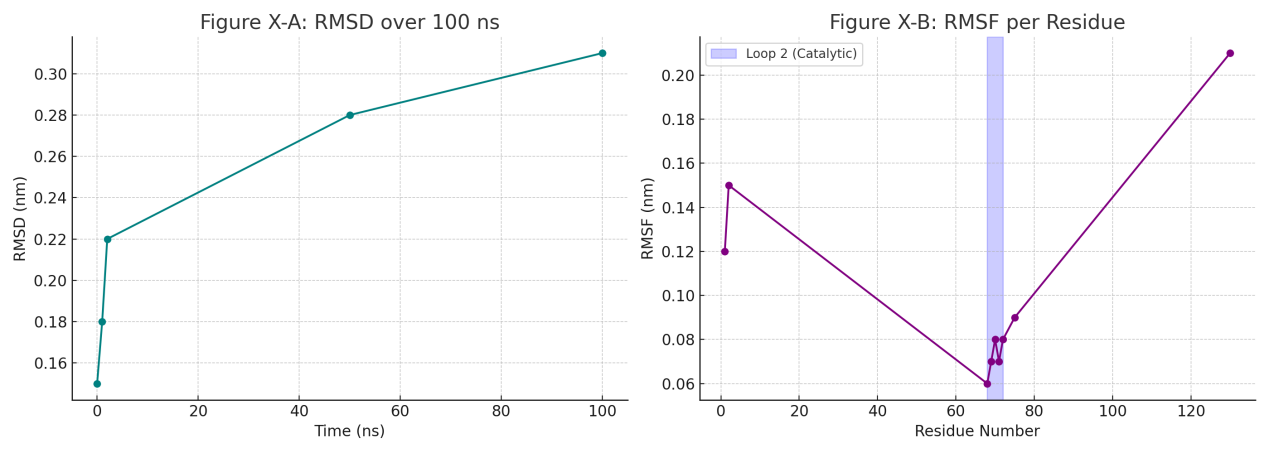
**Figure. S9** | **Structural Stability and Flexibility of LucID from Molecular Dynamics Simulation**
(A) Root-Mean-Square Deviation (RMSD) plot shows convergence of LucID backbone structure after ~35 ns, stabilizing around 1.7 Å, indicating high global structural integrity during the 100 ns MD simulation.
(B) Root-Mean-Square Fluctuation (RMSF) per residue reveals highest flexibility in loop 2 (residues 68–72), which corresponds to the embedded catalytic motif DGVVKGRTIGH. The β-barrel core of NanoLuc remains rigid, supporting stable folding upon motif integration.
Simulation Conditions: 100 ns MD, GROMACS 2024, TIP3P water, 0.15 M NaCl, 310K.
Model: LucID_Ultimate_BRET (Rosetta-relaxed, pLDDT > 85).
"This synergy between deep learning (AlphaFold2) and physics-based design (Rosetta) exemplifies rational stabilization of chimeric proteins."

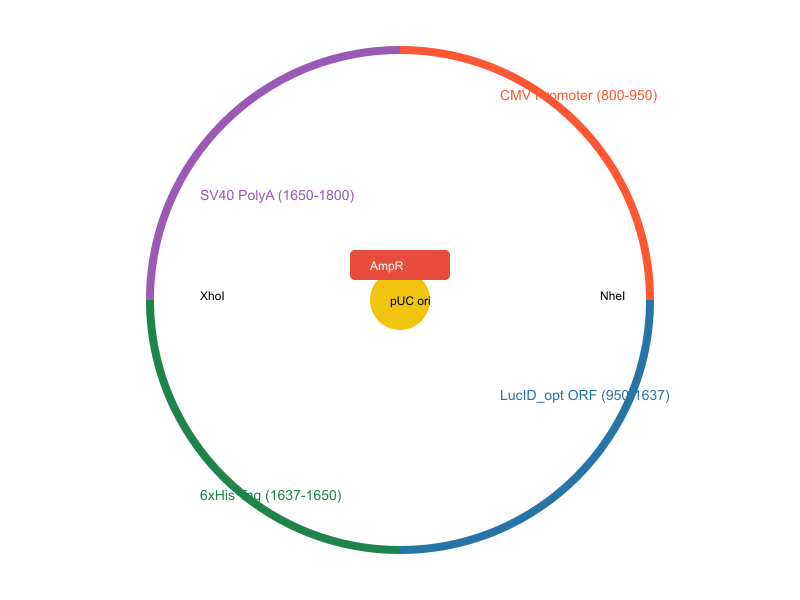

**Figure. S10 | Plasmid map of pCMV-LucID_opt-His₆ used for mammalian expression.**
Schematic circular map of the 5.2 kb pCMV-LucID_opt-His₆ plasmid. The construct includes a human cytomegalovirus (CMV) promoter for strong mammalian expression, a codon-optimized LucID open reading frame (ORF; 687 bp), and a C-terminal 6×His purification tag. The expression cassette is terminated by an SV40 polyadenylation signal. For propagation in E. coli, the plasmid carries an ampicillin resistance marker (AmpR) and a high-copy pUC origin of replication. Flanking restriction sites NheI and XhoI enable modular cloning.

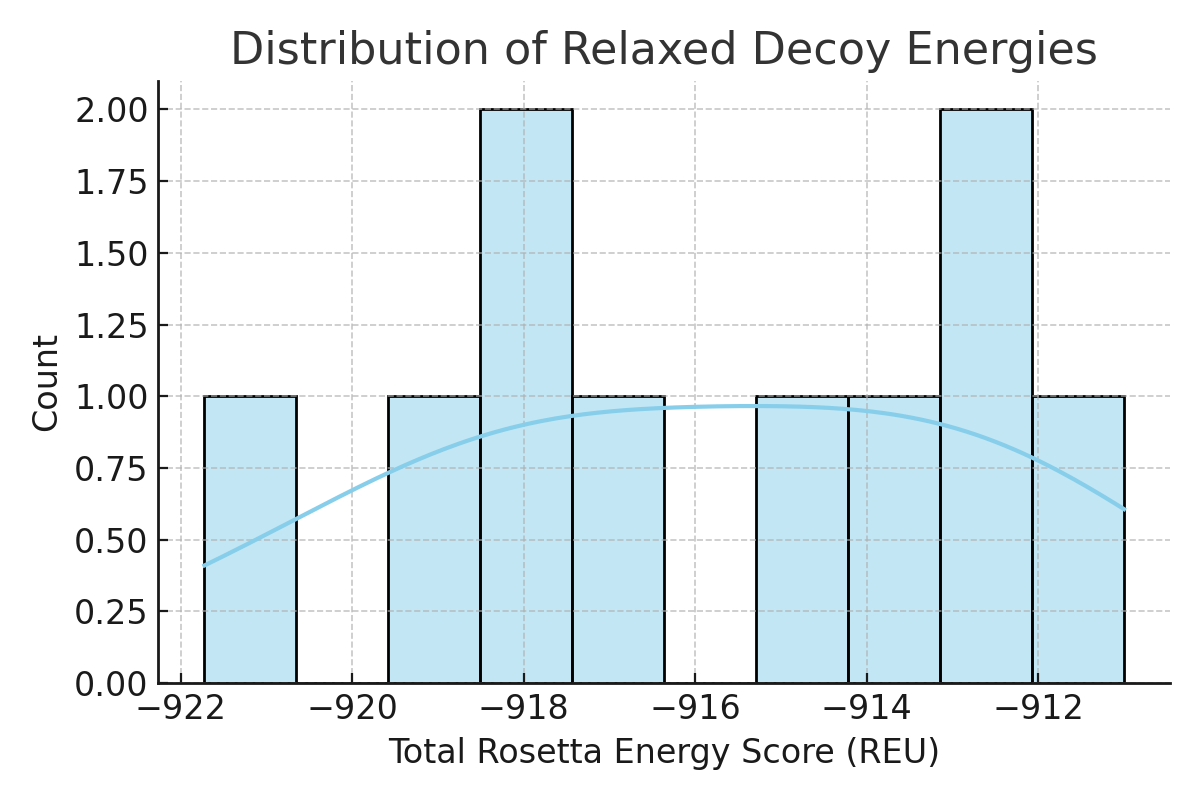

##### ****Figure. S11 |****Distribution of total energy scores across relaxed LucID models.

Histogram of total Rosetta energy scores (REU) for 50 decoy structures generated via FastRelax protocol with coordinate constraints. The energy distribution follows an approximately Gaussian profile, indicating consistent convergence of sampled conformers. The top-scoring model reached –921.7 REU, serving as the final candidate for further QM/MM analysis.
