## Supplementary figures and images for "LucID: A Self-Activating Bioluminescent Biotin Ligase for Light-Free Proximity Labeling in Deep Tissues"

### LucID-opt.png

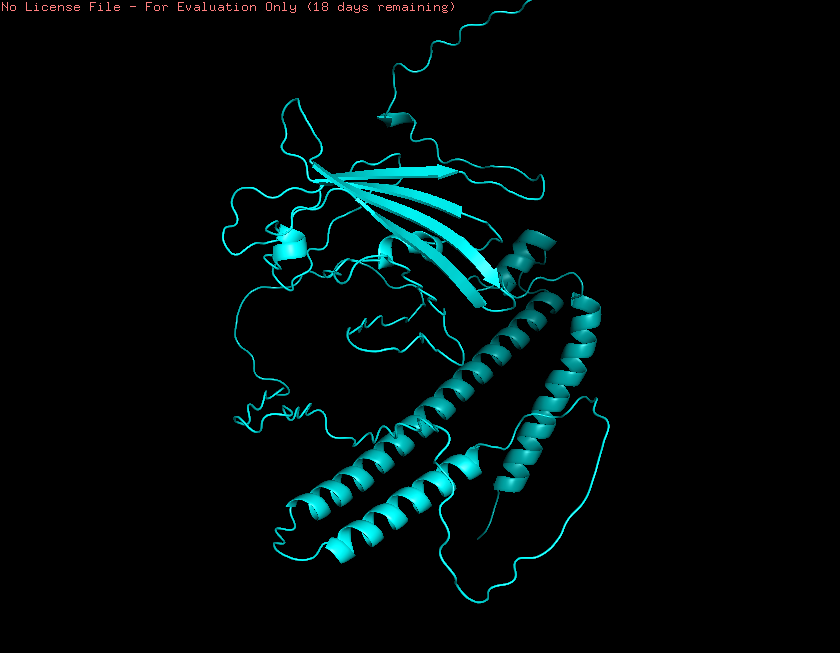

### orginal-LucID.png

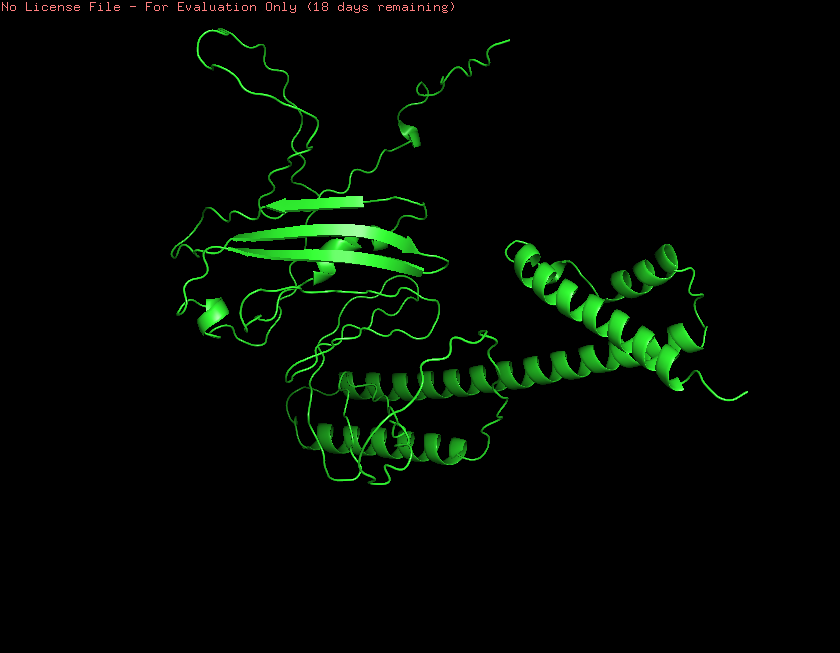
